# Functional relocation of the maize chloroplast *atpB* gene to the nucleus restores photosynthetic competence to a gene-edited non-photosynthetic mutant

**DOI:** 10.1101/2025.02.07.637139

**Authors:** Venkata RamanaRao Mangu, Kelsey Jenkins, Anna Pratt, Kara A Boltz, Leila Pazouki, Alice Y. Hui, Jeffrey M. Staub

## Abstract

We present a novel approach to photosynthetic gene engineering in maize using a nuclear-encoded, chloroplast-targeted TALE-cytidine deaminase enzyme to create non-photosynthetic knockout mutants of the chloroplast *rbcL* gene. An off-target mutation in the adjacent *atpB* gene, encoding the β subunit of ATP synthase, was consistently found in all edited lines, identified as pigment-deficient in tissue culture. These double mutants, carrying mutations in both genes, were purified to homoplasmy using unique leaf-base regeneration techniques. To test mutation complementation and identify the causal gene, nuclear transgenic lines expressing chloroplast-targeted RbcL and AtpB proteins were generated. The results show that nuclear expression of AtpB restores chlorophyll accumulation and supports wild-type growth in tissue culture.

---

Nonphotochemical quenching (NPQ) function was restored, and the maximum quantum yield of photosystem II (*Fv/Fm*) reached about 30% of wild-type levels in the nuclear transformed lines. This is the first demonstration in a monocot plant that complementation of a photosynthetic mutant via nuclear expression of the gene is possible, providing a facile method for future photosynthetic engineering.

The global population is projected to reach 9.6 billion by 2050, increasing food demand while available arable land shrinks and conventional breeding methods yield slow and limited productivity gains. This trend underscores the need for advanced bioengineering solutions to boost crop yields and improve resource efficiency. Enhancing photosynthetic capacity is a promising strategy, but most research focuses on C3 plants, despite the significant presence of C4 species in agriculture and their crucial role in providing food, feed, and fuel.^1–2^

Nuclear transgenic strategies have been used and multiple approaches proposed to enhance photosynthetic parameters to improve crop yields.^3–5^ In some cases, overexpression of endogenous or heterologous components of the nuclear-encoded electron transport machinery is associated with increased growth or enhanced photosynthetic parameters and recent examples for this approach show promise in some C4 plants. For example, in *Setaria* and sorghum, overexpression of the Rieske FeS subunit of the cytochrome *b_6_f* complex led to either an increased rate of photosynthetic electron transport or a faster response of non-photochemical quenching (NPQ), respectively, resulting in higher CO_2_ assimilation rates and biomass.^6–7^ In maize, research by Salesse-Smith et al.^8^ showed that the simultaneous overexpression of the Rubisco large and small subunits, along with the RAF1 assembly chaperone, increased Rubisco content by over 30%, enhanced CO_2_ assimilation rates by 15%, and improved fresh weight.

Surprisingly, photosynthesis can also be enhanced by nuclear overexpression of certain chloroplast-encoded proteins in the nucleus. For instance, the nuclear overexpression of the chloroplast *psbA* gene, which encodes the D1 protein of Photosystem II, has been shown to improve CO2 assimilation rates and increase biomass in plants such as *Arabidopsis*, tobacco, and rice.^9^ In a previous study, a nuclear-encoded, atrazine-resistant D1 protein from *Amaranth* was shown to successfully integrate into the thylakoid membranes of tobacco.^10^ Additionally, the nuclear expression of the chloroplast-encoded *rbcL* gene was able to compensate for a knockout of the gene in tobacco chloroplasts, achieved through plastid transformation technology.^11^

Plastid transformation technology has been extensively utilized in tobacco, where it is now routine for engineering chloroplast-encoded photosynthetic genes.^12–13^ Much of the research has concentrated on enhancing carboxylation properties of Rubisco by replacing the native *rbcL* gene in tobacco with heterologous *rbcL* genes from plants, bacteria, or algae (with or without the incorporation of small subunit genes to enhance Rubisco assembly).^14^ More recently, Rubisco engineering in potato via plastid transformation was reported, involving the replacement of the endogenous *rbcL* gene with the heterologous *Rhodospirillum rubrum* gene.^15^ Research has also examined components of the tobacco electron transport chain, with efforts including knockout studies^16^ and mutational analyses of chloroplast-encoded *atpB* subunit gene of the ATP synthase.^17–18^

The technology for stable chloroplast transformation in monocots has not yet been developed,^19^ preventing plastid transgenic approaches to photosynthesis research in commercially important C4 plants. On the other hand, recent advancements in organellar gene editing technologies^20–30^ have begun to enable engineering of chloroplast- and mitochondrial-encoded traits, including CMS^31–35^ and herbicide tolerance.^36–37^ Engineering of photosynthesis via editing of the chloroplast-encoded *rbcL* gene in *Arabidopsis* also appears possible.^38^

As an initial step towards a novel approach for photosynthetic gene engineering in maize, we utilized a nuclear-encoded, chloroplast-targeted TALE-cytidine deaminase enzyme to create a nonphotosynthetic knockout mutant of the maize chloroplast *rbcL* gene. Along with the *rbcL* loss of function, we identified an off-target mutation in the adjacent *atpB* gene, which complicated the analysis of photosynthetic parameters. Through nuclear expression of chloroplast-targeted RbcL and AtpB proteins, we identified AtpB as being responsible for a partial rescue of photosynthetic function in the double mutant lines. We demonstrated that nuclear-encoded AtpB fully restores NPQ and photosynthetic yield to about 30% of wild-type levels in transgenic plants under ambient conditions. This approach confirms that the nuclear expression of a chloroplast-encoded component of the photosynthetic machinery is functional in maize and offers a straightforward transgenic strategy to further explore the roles of chloroplast-encoded photosynthetic proteins.

## Results

We sought to use TALE-DddA-UGI cytidine deaminase^39^ base editing to knock out the maize chloroplast *rbcL* gene, encoding the large catalytic subunit of Rubisco. Maize nuclear codon-optimized transgenes (Fig. 1A) for the TALE-DddA-UGI left and right fusion proteins were each designed with a chloroplast transit peptide at their N-terminus for targeting of the proteins to the chloroplast compartment. The TALE-DddA-UGI transgenes were driven by maize or rice ubiquitin promoters for high-level expression and cloned into a T-DNA vector carrying a plant selectable marker gene (*bar*) encoding resistance to bialaphos and phosphinothricin, to create the pPTS424 vector. The pPTS424 vector was introduced via *Agrobacterium*-mediated transformation into immature maize embryos of a wild-type and a transgenic line overexpressing the morphogenic *Babyboom* (BBM) and *Wuschel2* (WUS2) genes (see below).

**Figure 1.**
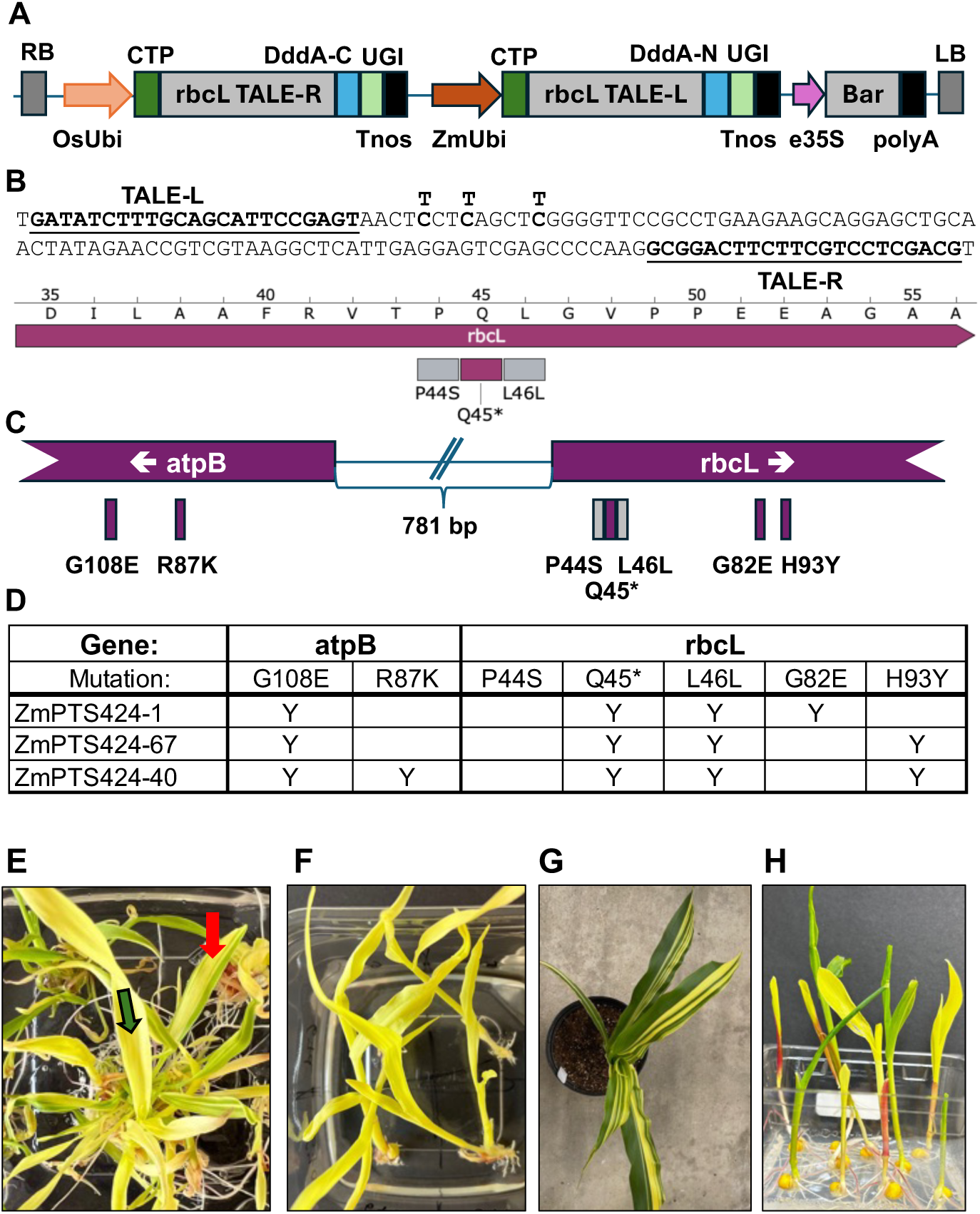
TALE-cytosine deaminase-mediated mutations in the maize chloroplast genome A) T-DNA region of vector pPTS424, carrying the “left” and “right” TALE-DddA-UGI nuclear-encoded chloroplast-targeted transgenes and the selectable *bar* gene. B) Targeted TALE binding sites in the *rbcL* gene coding region, showing the possible C to T edit sites and potential resulting amino acid changes. C) Compilation of the observed targeted *rbcL* gene mutations and off target mutations in *rbcL* and the adjacent *atpB* gene D) Summary of amino acids changes in *rbcL* and *atpB* genes in independent mutant lines. ZmPTS424-1 in is the wild-type genetic background whereas ZmPTS424-67 and ZmPTS424-40 lines are in the *BBM/WUS2* genetic background E) Phenotypic of T0 mutant plants after the initial plant regeneration. Note uniformly pigment deficient plantlets (green arrow) and plantlets with chimeric leaves (red arrow). F) Representative homoplasmic chloroplast mutant lines after two rounds of plant regeneration from leaf-base or nodal sections. G) Chimeric plants carrying green wild-type leaf sectors and mutant (yellow) leaf sectors growing in soil in the greenhouse. H) Segregation in T1 seedlings of pigment deficient homoplasmic chloroplast mutant and green wild-type seedlings.

Based on TALE-binding site requirements^40–41^ (Christian et al., 2010; Cermak et al., 2011) and DddA cytidine deaminase target site preference,^39^ a C to T base edit was targeted to create a premature stop codon at amino acid 45 of the *rbcL* N-terminus (Q45*), expected to completely abolish protein accumulation (Fig. 1B). The 22-nucleotide (nt) spacer region between 23 nt and 20 nt left and right TALE binding sites, respectively, also carries two additional potential C to T targets, at the amino acids immediately upstream and downstream of the intended stop codon mutation. C to T transversion of the upstream target could lead to a proline to serine amino acid change (P44S) while transversion of the downstream C to T target (L46L) would not result in an amino acid change (Fig. 1B).

Pigment deficient plants were previously obtained in T0 rice plants after editing of plastid-encoded photosynthetic genes,^30,42^ though these were not utilized further due to the lethality of those mutations in soil and the lack of a tissue culture system to support their growth. Although the phenotype of a maize plastid-encoded *rbcL* knockout is unknown, we anticipated a pigment deficient phenotype as has been observed in homoplasmic tobacco *rbcL* deletion plants^11^ or nuclear mutants in maize.^43^ To facilitate subsequent tissue culture maintenance and use of pigment mutant lines, we used as recipient for nuclear transformation a previously established maize line that overexpresses the morphogenic *BBM* and *WUS2* genes. In maize, embryogenic callus can be recovered from leaf-base tissue of transgenic *BBM/WUS* lines,^44^ while only rarely successful from some wild-type genotypes,^45^ thus enabling ongoing tissue culture maintenance of chloroplast-edited pigment deficient T0 plants.

Nuclear-transformed lines were recovered in each genotypic background after ∼6-8 week of bialaphos selection (Fig. 1C, D). Since chloroplast edits in other species were heteroplasmic in calli and T0 lines, we allowed nuclear transformed calli to regenerate into numerous plantlets so that pigment deficient leaf sectors or plants could be identified visually. In the first transformation experiment of each genetic background, a total of three independently transformed events that regenerated pigment deficient plants were further studied (Fig 1D). In each line, regenerated plants carried either large pigment deficient leaf sectors or entirely pigment deficient plants (Fig. 1E), indicating that a majority of the plastid genomes in each plant carried edits prior to plant regeneration.

### Purification of pigment deficient lines

To enable long-term study of pigment deficient mutants, it was important to ensure homoplasmy and stability of plastid genome edits in the mutant lines. In tobacco chloroplast transformation experiments, multiple rounds of regeneration from leaf cells of transplastomic lines is used to routinely achieve homoplasmy.^46^ In most monocots such as maize, regeneration from leaf cells is typically not possible. However, the *BBM/WUS2* overexpression genetic background in some of our mutant lines could enable efficient recovery of regenerable embryogenic callus from leaf base tissues.^44^ Using the basal ∼1 cm of leaf-base tissue from mutant line ZmPTS424-67 plants that initially appeared uniformly pigment deficient, small dissected basal leaf pieces were placed onto high auxin media where embryogenic callus formed at high-efficiency from nearly every tissue piece within 10-14 days.

Regenerable embryogenic callus could also be recovered from nodal segments of the ZmPTS424-1 mutant line in the wild-type genetic background, albeit at much lower efficiency than in the *BBM/WUS2* lines. In this case, the meristematic region embedded within the first nodal region of young pigment deficient plants grown in vitro was dissected and placed onto high auxin media.^47^ Embryogenic callus formed sparingly within ∼3 weeks, however, once established, embryogenic callus was prolific and grew nearly as well as that from the ZmPTS424-67 line. Embryogenic callus from mutant lines in both genetic backgrounds were then routinely used to re-regenerate numerous plants in the light to establish complete penetrance of the pigment deficient phenotype in both mutant lines (Fig. 1F). Using this approach, mutant plant lines were subjected to two rounds of additional plant regeneration to ensure homoplasmy of the multiple mutations in each plant line (see below).

Although uniformly pigment deficient plants could not survive in soil, as a second approach to obtain uniformly pigment deficient and homoplasmic mutant lines, we grew chimeric ZmPTS424-1 plants carrying a mixture of green and pigment deficient sectors in soil in the greenhouse to maturity (Fig. 1G, H). T1 seeds from these lines were expected to segregate uniformly green, uniformly pigment deficient or chimeric T1 seedlings if the plastid mutations were germline transmitted. Moreover, pollen from wild-type plants was used to fertilize the plastid mutant plants to facilitate recovery of seedlings that no longer carry the TALE-DddA-UGI nuclear transgene, eliminating the possibility of acquiring additional mutations during long-term growth of mutant lines. As can be seen in Fig. 1I, uniformly pigment deficient T1 seedlings were recovered along with chimeric plants that were replanted in the greenhouse for mutant seed amplification, providing a source of genetically stable material for ongoing experiments. PCR analysis was also used to identify segregated T1 seedlings that no longer carry the nuclear TALE-DddA-UGI transgenes (Supplementary Table 1) ensuring stability of the mutant lines.

### Pigment deficient lines carry homoplasmic chloroplast genome mutations in adjacent photosynthetic genes

PCR-sequencing of the plastid *rbcL* genomic region was used to confirm editing of the intended target sites during rounds of plant regeneration and in T1 seedlings, while chloroplast genome sequencing was used to confirm homoplasmy of TALE-DddA-UGI-derived mutations and screen for potential plastid genome off-target edits. As shown in Fig. 1 C, D, each of the mutant lines carried based edits only in the *rbcL* gene and the adjacent *atpB* gene.

The intended *rbcL* Q45* stop codon edit and the adjacent silent L53L edit were recovered together in all lines while the P44S edit did not occur in any line (Fig. 1D). Interestingly, the unedited P44S site is in a 5’-TC context previously shown to be favored by the fusion enzyme,^39^ though its close proximity to the left TALE binding site may have created a steric hindrance that prevented editing. Off-target C to T edits resulting in amino acid changes were also recovered downstream in the *rbcL* coding region of mutant lines ZmPTS424-1 and ZmPTS424-67, though these are inconsequential relative to the Q45* stop codon. Mutant line ZmPTS424-40 had the same edits observed in the previous mutant lines and an additional C to T edit creating R87K of the *atpB* gene. However, during the second re-regeneration of plants from this line, multiple additional chloroplast genome mutations occurred in several locations outside of the *rbcL* and *atpB* genes that made further work with this line not feasible.

Whole genome sequencing of twice-re-regenerated mutant plants summarized in Table 1 indicated that lines ZmPTS424-1 and ZmPTS424-67 carried homoplasmic plastid genome edits described above with no other off-target edits, and were used for all subsequent studies. Furthermore, periodic plastid genome sequencing has been repeated multiple times over more than 1.5 years of tissue culture maintenance with similar results, indicating that these lines are stable.

### Nuclear complementation of the plastid-encoded pigment deficiency

Since all pigment mutant lines carried the Q45* stop codon knockout of *rbcL* and the G108E mutation in the adjacent *atpB* gene coding region, it was critical to determine which mutated gene caused the pigment deficiency. In tobacco and maize, mutation in either gene causes pigment deficiencies with different severity.^11,17–18,48^ Since chloroplast transformation technology is not available in maize, we sought to identify the causative gene and complement the chloroplast mutants via nuclear expression of the wild-type chloroplast photosynthetic genes by re-targeting of the encoded proteins back to the chloroplast.

T-DNA vectors were constructed that carry maize codon-optimized wild-type *rbcL* (pPTS454) or *atpB* (pPTS451) coding region, translationally fused at the N-terminus to a chloroplast transit peptide and at the C-terminus to an epitope tag for subsequent detection of the transgenic recombinant proteins and discrimination from the cognate endogenous chloroplast-encoded proteins (Fig. 2A). Although maize Rubisco is only functional in bundle sheath cells,^49^ we expressed the transgenes from strong constitutive promoters to coordinate expression of the two proteins and to facilitate high-level expression that might be needed for complementation of a chloroplast mutation. Nuclear retransformation via particle bombardment of embryogenic callus derived from the homoplasmic ZmPTS454-1 and ZMPTS454-67 mutant lines was performed using selection for a hygromycin resistance transgene on the T-DNA vectors. Cotransformation of the two T-DNA vectors was used to facilitate recovery of lines expressing either *CTP-rbcL*, *CTP-atpB*, or both *CTP-rbcL* and *CTP-atpB* transgenes.

**Figure 2.**
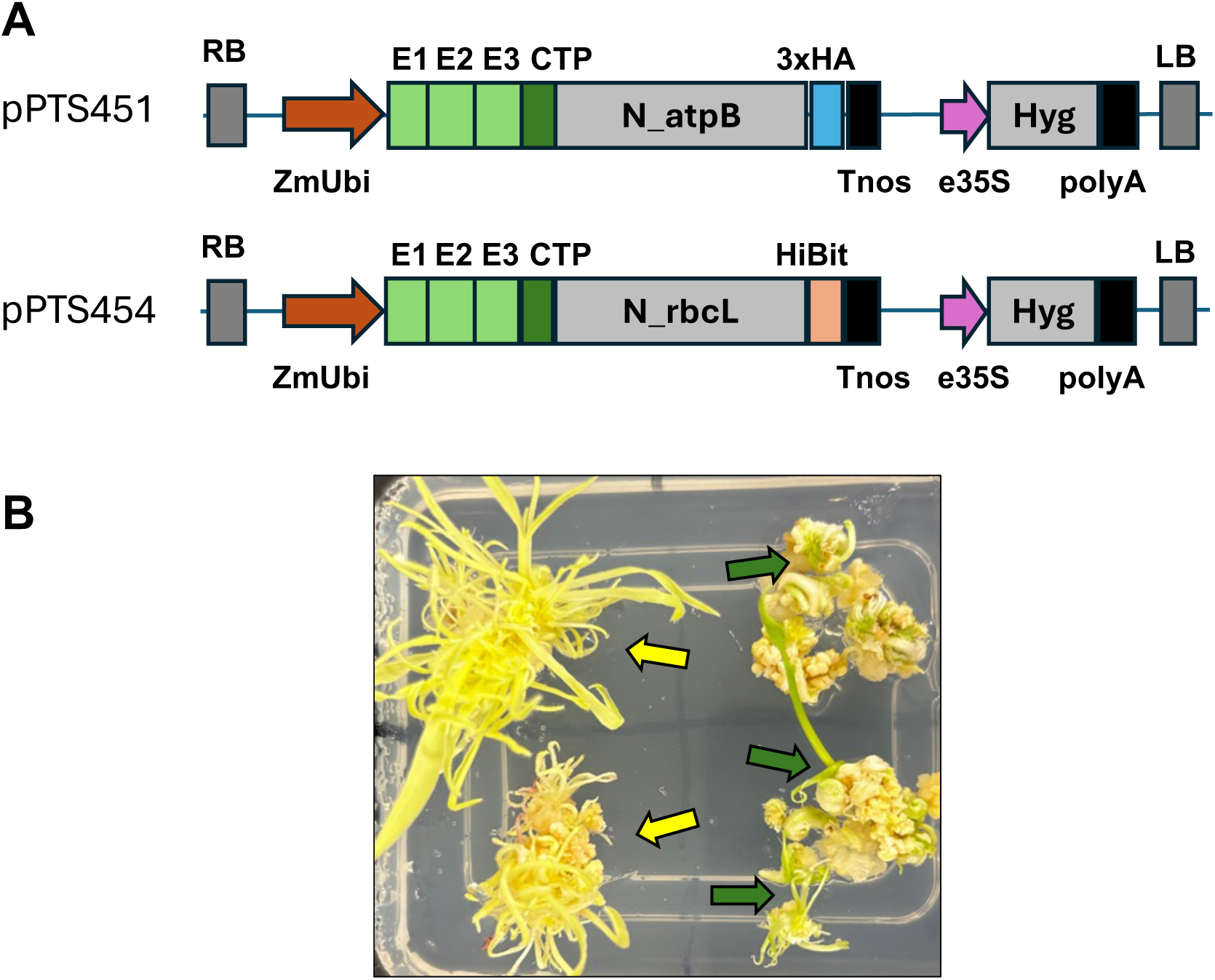
Retransformation of homoplasmic chloroplast mutant lines with nuclear-encoded chloroplast-targeted *atpB* or *rbcL* transgenes. A) T-DNA regions of pPTS451 and pPTS454 binary vectors carrying maize codon-optimized *atpB* or *rbcL* transgenes, respectively. Both coding regions are translationally fused at the N-terminus to the Rubisco small subunit 2 gene chloroplast transit peptide (CTP) and at the C-terminus by either a 3xHA epitope (pPTS451) or HiBit epitope tag (pPTS454) for discrimination in plants. Expression of the transgenes is driven by three viral enhancers (E1, E2, E3) and the maize ubiquitin 1 gene promoter (ZmUbi), and a nos terminator (Tnos). The selectable hygromycin resistance gene is driven by enhanced CaMV 35S (e35S) promoter and polyA signal. B) Hygromycin resistant nuclear re-transformation of the chloroplast mutant lines with the pPTS451 and pPTS454 vectors results in either regenerating pigment deficient lines (yellow arrows) or green complemented plantlets (green arrows).

Hygromycin resistant callus lines were regenerated in the light to look for green plantlets that would indicate complementation of the pigment deficient phenotype of the mutants. Multiple independently retransformed T0 events regenerated as green plants indicating complementation of the mutant phenotype while other lines regenerated only pigment deficient lines indistinguishable from the parental lines (Fig. 2B). PCR analysis was used to verify the presence or absence of the recombinant *CTP-rbcL* and/or *CTP-atpB* transgenes in the lines (Supplementary Table 2) to identify lines with either a single transgenes or both transgenes. Based on the PCR results, the presence of the *CTP-atpB* transgene strictly correlated with green complemented lines, while *CTP-rbcL* did not, indicating that the pigment deficiency was likely due to the G108E edited chloroplast *atpB* gene in the mutant lines. The complementation results were confirmed by transformation of the pPTS454 *CTP-rbcL* or pPTS451 *CTP-atpB* T-DNA vectors separately, which verified complementation of the pigment deficiency via the *CTP-atpB* transgene only (data not shown). PCR-sequencing confirmed the original chloroplast *rbcL* and *atpB* gene mutations were still present in all nuclear complemented lines, ruling out any reversion of the original mutations (Supplementary Table 2).

### Nuclear transgenic ATPB protein expression is sufficient to rescue the pigment deficient phenotype

While the presence of the *CTP-atpB* transgene correlated with green plants in retransformed lines, it was important to verify that nuclear-encoded AtpB protein expression also correlated with the complemented phenotype. Protein blot analysis (Fig. 3) was used to distinguish recombinant AtpB protein via its HA-epitope tag from any potential endogenous chloroplast-encoded AtpB protein, which was detected via polyclonal antibody. Fig. 3A shows that wild-type green plants accumulate abundant AtpB protein, as expected, while the mutant parental lines ZmPTS424-1 and ZmPTS424-67 show only weak hybridization with the polyclonal antibody. Since the polyclonal antibody to AtpB also recognizes the mitochondrial-encoded subunit, it’s unclear whether the weak hybridization signals observed in the mutant lines represent residual chloroplast-encoded ATPB Q108E protein, mitochondrial AtpB, or both. In contrast, leaf protein extracts from every nuclear retransformed line (designated as 451+4 followed by the event number in Fig. 3) that carries the *CTP-atpB* transgene and is green also accumulates recombinant AtpB protein, as revealed by hybridization with the HA-epitope tag antibody (Fig. 3A middle). The slightly different mobility in the protein gel blot of the HA-tagged AtpB protein clearly distinguished it from any endogenous protein (Fig. 3C). In contrast, none of the retransformed lines that remained pigment deficient accumulated any detectable recombinant AtpB protein.

**Figure 3.**
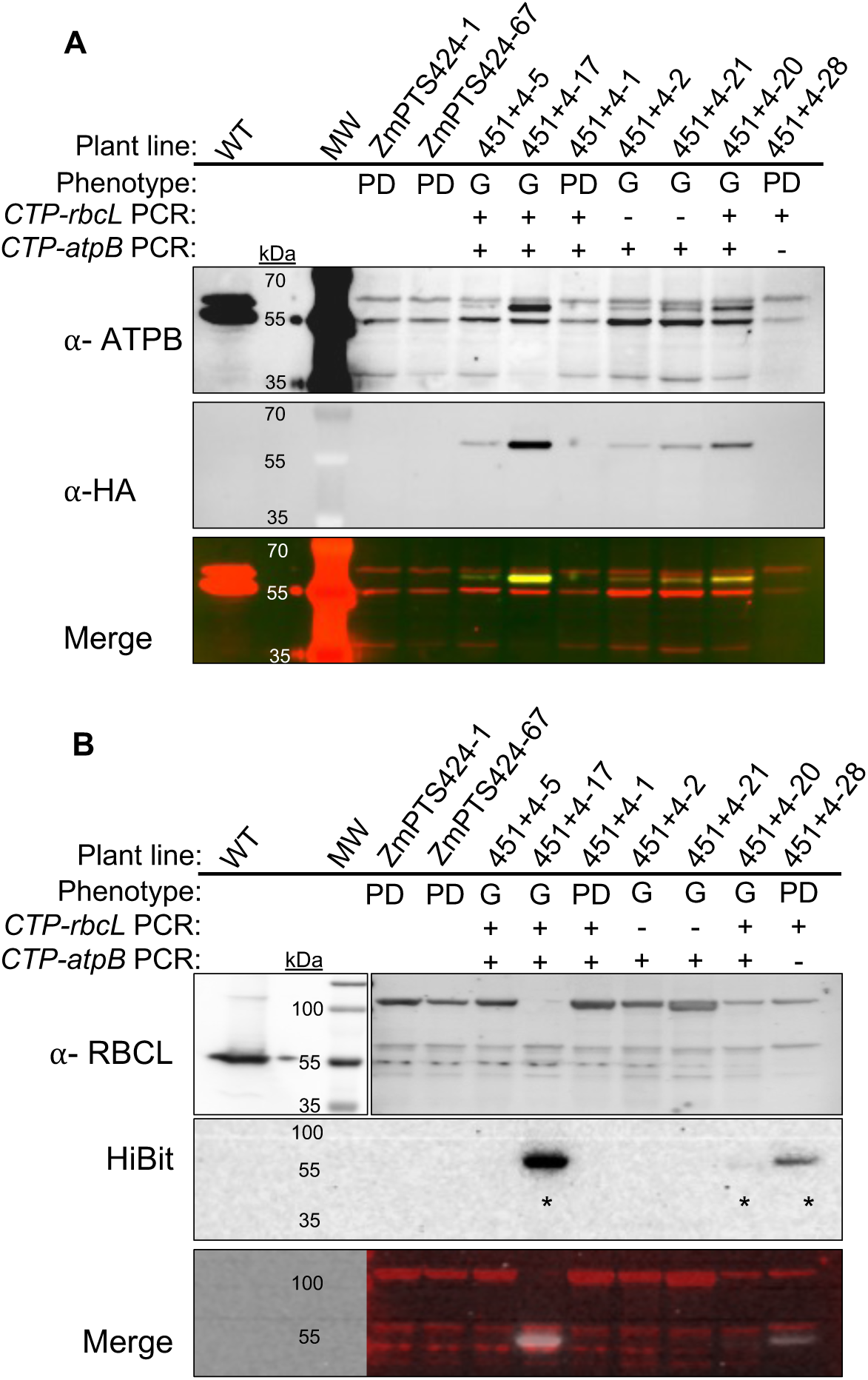
Protein gel blots of nuclear re-transformed chloroplast mutant lines. For both blots, the protein amounts loaded were 20 μg for wild type and 40 μg for all other samples. The phenotype (PD = pigment deficient; G = green) and PCR-based genotype result (+ or – to indicate presence or absence of the nuclear complementation construct) are indicated for each sample. The chloroplast mutant lines ZmPTS424-1 and ZmPTS424-67 are included, as well as nuclear retransformed lines which are designated as 451+4 followed by the event number. (A) Detection of total AtpB in leaf protein extracts using α-AtpB antibody (top) and transgenic AtpB using α-HA epitope tag antibody (middle). The bottom panel shows the merge of the two signals. AtpB-HA protein is easily observed in the top and merge blots due to its unique molecular weight. (B) Detection of total RbcL protein in the same leaf protein extracts using α-RbcL antibody (top) and transgenic RbcL protein using luminescent detection of the HiBit epitope tag (middle). The bottom panel shows the merge of the two signals. A short exposure of the α-RbcL antibody blot is shown (separate box on left) for wild-type (WT) leaf extract to enable easy viewing of the correct molecular weight of RbcL. * Samples with RbcL-HiBit luminescent signal. MW = molecular weight markers. Original, uncropped images are available in the Supplemental materials.

Protein gel blot detection of chloroplast-encoded and recombinant RbcL was also performed (Fig B). Polyclonal antibody shows the wild-type accumulates highly abundant RbcL protein as expected, while the parental mutant lines do not accumulate any detectable chloroplast-encoded protein, though several apparent non-specific bands are present in all samples (Fig. 3B top). Nuclear-encoded recombinant RbcL protein, distinguished by the HiBit epitope tag (Fig. 3B middle), does accumulate in some green and non-green lines, confirming lack of correlation to the complementation phenotype.

As expected for nuclear transformation events, each retransformed line carries a different amount of recombinant AtpB or RbcL protein. In green complemented lines, recombinant AtpB protein accumulates at dramatically lower levels than the wild-type AtpB protein. Using a dilution series of the wild-type protein extract, we estimate that recombinant AtpB accumulates up to only ∼5% of the native chloroplast-encoded protein (Fig. 4). Surprisingly, this lower level of AtpB protein is apparently enough to enable chlorophyll accumulation and photosynthesis in complemented lines.

**Figure 4.**
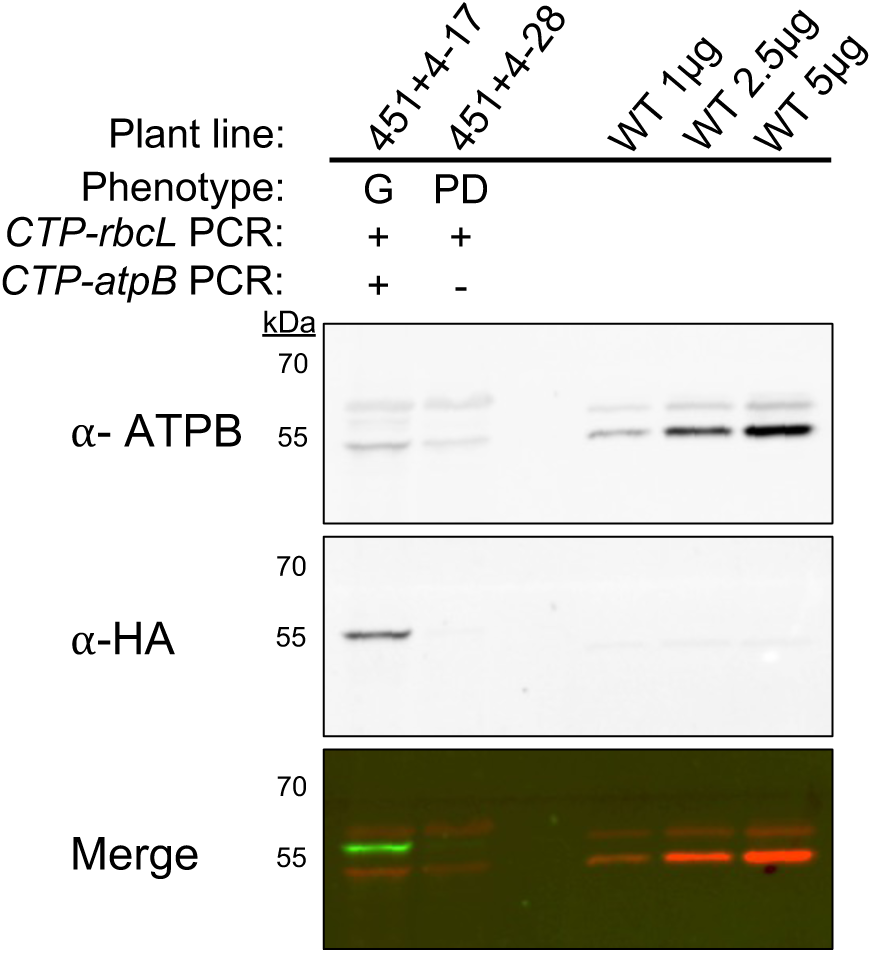
Comparison of AtpB protein levels in nuclear-retransformed lines and wild type plants. 40 μg of total leaf protein was loaded on the protein gel blot for the representative 451+4-28 pigment deficient (PD) and green (G) complemented 451+4-17 nuclear retransformed lines. Wild-type leaf protein extract was loaded at 1 ug, 2.5 ug, and 5 μg to enable estimated quantification of AtpB protein level in the complemented line. The genotype (+ or – to indicate presence or absence of the nuclear complementation construct) are indicated for each nuclear retransformed line. Original, uncropped images are available in the Supplemental materials.

### Rescue of photosynthetic function in complemented lines

The pigment deficient phenotype of chloroplast mutants and lack of AtpB and RbcL proteins clearly shows impairment of photosynthetic activity whereas the green phenotype and correlated accumulation of transgenic AtpB protein in complemented plants points to significant improvements in the affected photosynthetic parameters. Since the parental chloroplast mutant line has primary lesions in both the Calvin cycle due to loss of Rubisco and a disruption in proton motive force (pmf) across the thylakoid membrane due to loss of ATPase synthase, it was important to gain insight into which photosynthetic parameters are restored in the nuclear complemented lines. As shown in Fig. 5, *Fv/Fm* (the maximum quantum yield of Photosystem II) was determined by chlorophyll fluorescence of dark-adapted plants subjected a saturating pulse of high actinic light. NPQ, the photoprotective mechanism that helps dissipate excess light energy triggered by the change in pH across the thylakoid membrane, was also determined.

**Figure 5.**
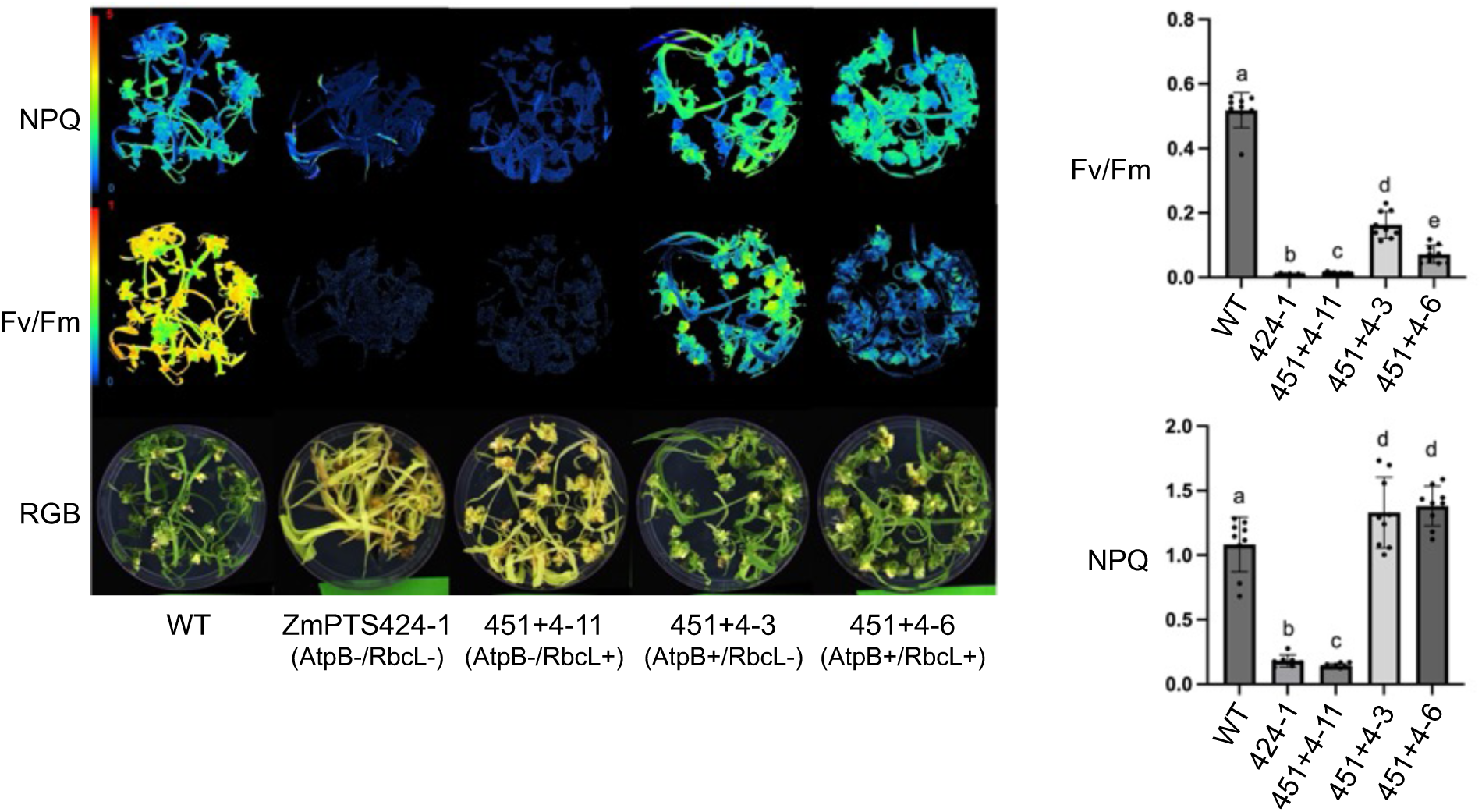
Measurements of photosynthetic parameters *F_v_/F_m_* and NPQ. Plantlets were dark adapted for 20 minutes and then subjected to a pulse of saturating light, and then a light induction curve. A) NPQ and *F_v_/F_m_* images are false colored images to illustrate chlorophyll fluorescence differences. B) *F_v_/F_m_* and C) NPQ values are shown graphically. Lower case letters indicate values with different statistical significance, as measured by Student’s T-test. The number of plantlets for each genotype is as follows: WT = 13; chloroplast mutant line ZmPTS424-1 (AtpB-/RbcL-) = 6; nuclear retransformed lines 451+4-11 (AtpB-/RbcL+) = 14; 451+4-3 (AtpB+/RbcL-) = 12; 451+4-6 (AtpB+/RbcL+) = 15.

As shown in Fig. 5A, callus cultures from each plant line were allowed to regenerate into plantlets for ∼3-4 weeks to synchronize growth of all lines to the same stage prior to chlorophyll fluorescence analysis. In addition to wild-type and parental ZmPTS424-1 mutant plants, representative nuclear transgenic lines expressing different combinations of AtpB and RbcL were tested, including pigment deficient (451+4-11, AtpB-/RbcL+), and green complemented lines (451+4-3, AtpB+/RbcL- and 451+4-6, AtpB+/RbcL+). Not surprisingly, chlorophyll fluorescence images indicate little or no photosynthetic activity (measured by *Fv/Fm* and NPQ) under these conditions in the parental mutant line and the nuclear transgenic line that lacks AtpB (451+4-11). On the other hand, partial restoration of the photosynthetic function in complemented lines (451+4-3 and 451+4-6) that accumulate the AtpB protein is accompanied by significant increase in *Fv/Fm*, and NPQ measurements that appear similar to the wild-type control.

The quantitative data for *Fv/Fm* (Fig. 5B) and NPQ (Fig. 5C) confirm the significant restoration of photosynthetic activity in the complemented 451+4-3 and 451+4-6 lines. *Fv/Fm* levels in these lines (Fig. 5B) are restored to approximately 15–30% of wild-type levels, while NPQ values (Fig. 5C) are slightly higher than those of wild-type plants, showing a statistically significant increase. In contrast, the parental mutant and lines lacking AtpB have only basal levels of either photosynthetic parameter. These results clearly indicate that the nuclear expressed AtpB is imported into the chloroplast, incorporated into ATP synthase complex and is active in photosynthesis. On the other hand, the nuclear expressed RbcL protein appears not to contribute to the rescue of greening or photosynthetic parameters in these lines.

## Discussion

Plastid transformation technology has been available in multiple plants species for more than 25 years^12–13^ and has been used to study numerous aspects of plastid biology and to introduce potentially commercial traits, but stable plastid transformation technology has not been developed in any monocot crop. More recently, nuclear encoded gene editing tools have been used to modify organellar genes but have been limited to proof-of-concept reports that recover pigment deficient or antibiotic resistant lines as screenable markers for editing events. In limited cases, gene editing tools have generated potentially useful traits such as creating or curing plants of cytoplasmic male sterility^31–35^ or generating herbicide resistant plants.^36–37^ We have taken a different approach, to create non-photosynthetic mutants of maize and utilizing those to study the function of chloroplast genes via the much more facile approach of nuclear transformation. We have shown for the first time in a monocot plant that complementation of a photosynthetic mutant via nuclear expression of the gene and retargeting of the protein to the chloroplast is possible and a viable approach to enable easier study of photosynthetic parameters. A previous approach in tobacco^11^ utilized deletion of the *rbcL* gene through existing chloroplast transformation technology and subsequent relocation of the plastid gene to the nucleus, though no effort was reported to test photosynthetic parameters in those plants. Our approach reported here serves as a novel approach to begin the design and testing of nuclear-relocated chloroplast genes in plant species where plastid transformation technology does not yet exist.

The selection and maintenance of homoplasmic non-photosynthetic chloroplast mutants created via nuclear gene editing tools is complicated by the chloroplast polyploid genetic system and the lack of methods to segregate mutant genomes in monocot tissue culture. In tobacco, the standard approach to obtain homoplasmy utilizes a chloroplast-localized selectable maker and multiple regeneration steps from transformed leaf tissue,^14–15^ that is difficult or not possible in maize. We used multiple approaches to overcome these issues, including the use of overexpressed morphogenic genes to facilitate rapid recovery of callus from leaf base tissues, optimized callus induction media in the wild-type genetic background, and the recovery of chimeric plants that generated segregated homoplasmic mutants in T1 seed. Pigment deficient lines were easily identified during the selection phase for nuclear transgenic TALE-DddA-UGI events, suggesting that the TALE-cytosine deaminase derived mutations were initially very penetrant and amplified to multiple plastid genomes in the majority of cells of initial T0 regenerated plants. However, each of the mutant lines became homoplasmic for all mutations only after rescue of callus from leaf base or nodal sections of plants regenerated in tissue culture, as shown by deep chloroplast genome sequencing (Table I). Our ability to maintain homoplasmic mutant and nuclear complemented lines as callus cultures and to synchronize regeneration of plants at any time also facilitated the analysis of growth conditions and photosynthetic parameters in this study, which was not attempted in the only other reports of chloroplast gene editing in a monocot.^30,42^

**Table I.** Homoplasmy of chloroplast mutant lines determined by deep genome sequencing of twice-regenerated T0 plants. The two studied chloroplast mutant lines ZmPTS424-1 and ZmPTS424-67 with their C to T mutations and protein coding effects are listed, along with sequence read depth and mutation frequency. Note that variant frequencies may be less than 100% due to sequence and read mapping errors, and the presence of nuclear-localized chloroplast sequences. Mutations were found to be at similar levels in multiple sequencing runs, including in T0 plants and T1 seedlings, indicating homoplasmy.

| Mutant Line | Gene | Nucleotide Change | Amino Acid Change | Protein Effect | Read Coverage | Variant Frequency |
| --- | --- | --- | --- | --- | --- | --- |
| ZmPTS424-67 | *atpB* | C -> T | G108E | Substitution | 339 | 99.40% |
|  | *rbcL* | C -> T | Q45* | Truncation | 392 | 93.90% |
|  | *rbcL* | C -> T | L46L | None | 393 | 93.40% |
|  | *rbcL* | C -> T | H93Y | Substitution | 363 | 92.80% |
| ZmPTS424-1 | *atpB* | C -> T | G108E | Substitution | 359 | 98.60% |
|  | *rbcL* | C -> T | Q45* | Truncation | 400 | 93.80% |
|  | *rbcL* | C -> T | L46L | None | 411 | 93.40% |
|  | *rbcL* | C -> T | G89E | Substitution | 407 | 95.30% |

Chloroplast gene editing in our hands was efficient enough to recover pigment deficient lines in multiple nuclear transformation experiments using the pPTS424 TALE-DddA-UGI construct (Suppl. Table III). The intended *rbcL* stop codon target mutation was recovered in all characterized lines, indicating efficient design, chloroplast targeting and activity of the transgenic construct. The location of the stop codon mutation was chosen to simultaneously test editing efficiency at two additional potential C to T editing sites immediately adjacent. Interestingly, the silent L46L mutation occurred in each line but the P44S mutation did not occur in any line. Presumably, steric hindrance of the left TALE binding site prevented editing of the P44S site.

Notably, off-target mutations in both the *rbcL* and *atpB* genes were recovered in every transformed event and the identical off-targets were observed in multiple independent transformants, although no plastid genome sequences with obvious identity to the TALE binding site sequences are observed near these secondary mutations (Suppl. Fig. 1). Each of the mutations were stable in most mutant lines over numerous tissue culture subcultures, leaf regeneration cycles and through T1 seed generation. These results suggest that the either the TALE binding in those stable lines are remarkably specific and do not create new mutations during prolonged tissue culture, or we speculate that the TALE-DddA-UGI transgenes may have lost activity after the initial transformation event due to transgene silencing or some other mechanism. In contrast, one line (ZmPTS424-40, Fig. 1) was stable through two cycles of leaf base regeneration but subsequently accumulated multiple new mutations during tissue culture that precluded its use in further studies.

Deletion of chloroplast-encoded *rbcL* in tobacco^11^ or nuclear mutants in maize defective in Rubisco accumulation^43^ have a pale-green pigment deficient phenotype. On the other hand, maize nuclear mutations that affect chloroplast ATP synthase expression may be pale green^48^ whereas severe mutations in tobacco nuclear- or chloroplast-encoded ATP synthase subunits have an albino phenotype.^17–18^ Our nuclear complementation experiments confirm that the parental chloroplast *atpB* gene mutation causes a severe pigment deficient phenotype in maize, while we infer that a mutation in *rbcL* alone would have subtle or no apparent phenotype in tissue culture and therefore was not found in our mutant screen approach.

Analysis of numerous nuclear retransformed lines (Fig. 3 and Suppl. Table II) shows that green complemented lines carry the *CTP-atpB* transgene and accumulate the transgenic AtpB protein. Interestingly, AtpB protein accumulation was only up to ∼5% of wild-type protein levels in the highest nuclear expressing line studied (Fig. 4). The restoration of greening and photosynthetic activity indicate that the nuclear-encoded AtpB protein is correctly assembled with the other ATP synthase subunits and is properly localized in the thylakoid membrane complex. Previous work in tobacco has hypothesized that a large pool of inactive AtpB synthase content can be fully compensated for by the activation of a previously inactive enzyme population17 (Rott et al. 2011; Schottler et al., 2015), suggesting this may also be the case in maize. In contrast, RbcL protein accumulated at low levels in some plant lines (Fig. 3) but was not required for greening. Plants expressing AtpB were maintained in tissue culture, regenerated plants from callus, and rooted in tissue culture similarly to wild-type plants. However, neither AtpB+ or AtpB+/RbcL+ lines grew for longer than ∼2 weeks when transferred to soil in the greenhouse. This latter result may indicate that higher accumulation levels of AtpB and/or RbcL protein are required to support ongoing photosynthesis in the absence of the sugar typically added to tissue culture media. However, we did not attempt to grow plants under low light or other conditions that may have facilitated survival of complemented plants in soil. Overexpression of Rubisco with its assembly factor RAF^8^ did increase photosynthetic parameters in maize, indicating opportunities for potential improvement to our current approach.

We used chlorophyll fluorescence analysis to gain insight into the photosynthetic parameters affected in the parental mutant and nuclear transgenic lines. Plants were grown on sucrose-containing culture media to facilitate synchronized growth of regenerated shoots. Although tissue culture is known to reduce the overall rate of photosynthesis,^50^ wild-type plants have a high photosynthetic activity as measured by *Fv/Fm* under the conditions used (Figure 5A, B). In contrast, the parental mutant AtpB-/RbcL- and non-complemented transgenic AtpB-/RbcL+ line shows low levels of photosynthesis or barely detectable NPQ response, as would be predicted for plant lines that lack both electron transport and Calvin cycle functions. Interestingly, green complemented plants exhibit up to ∼30% of wild-type levels of *Fv/Fm* indicating significant restoration of Photosystem II activity based only on the presence of only ∼5% of wild-type AtpB protein levels and no apparent Rubisco activity. We infer that linear electron flow in complemented lines may have been partially restored via a Calvin cycle-independent mechanism such as the malate shuttle or other pathway,^52–53^ allowing some Photosystem II function in those lines. Importantly, NPQ function was restored, with the AtpB+ complemented lines showing elevated levels (Fig. 5C). This restoration of NPQ function may be due to a reinstatement of the ΔpH parameter of pmf through cyclic electron flow,^53^ which is not possible in the double mutant lines. The combination of both reduced ATP synthase activity and linear electron flow could result in an overacidification of the thylakoid lumen, triggering an elevated NPQ response.^54^

ATP synthase activity was abolished by a single amino acid change (G108E, Fig. 1) in the AtpB protein. That region of the protein is responsible for nucleotide coordination in the F1 ATPase multiprotein complex,^55–56^ and G108 is an invariant position as suggested by amino acid alignment across all available sequences in the NCBI database. AlphaFold structure prediction^57–58^ (www.alphafold.com; Supplementary Figure 2) suggests that G108 coordinates with Valine 232 (V232) to help maintain the structure of that ADP+ and Pi nucleotide binding region required for production of ATP. Since complemented transgenic lines incorporate nuclear-expressed chloroplast-targeted AtpB protein into the F1 ATPase, the transit peptide, which should be removed during protein import, and C-terminal epitope tag on the recombinant protein do not apparently disrupt appropriate assembly or activity in the complex.

These results indicate that our nuclear engineering approach can be used to easily design and test the function of AtpB variants in future work, including in response to factors such as light or drought stress, redox potential or phosphorylation state, sugar sensing and the assembly with other components of the enzyme complex.^54^ Other reports have begun to explore the role of specific regions of AtpB protein on function in the dicot plant, tobacco, using the existing chloroplast transformation technology.^17,59^ Moreover, future work will determine if nuclear-expressed AtpB also contributes to improvements in photosynthetic parameters in a wild-type genetic background, as has been reported for nuclear-encoded chloroplast *psbA* in tobacco, *Arabidopsis* and rice.^9–10,60^

## METHODS

### Vector construction

The left and right TALE-DddA-UGI^39^ transgene cassettes were assembled from synthetic DNA fragments on separate plasmids, then cloned next to each other to create the pPTS424 T-DNA transformation vector. All transgene protein coding regions were codon optimized for maize nuclear expression using Integrated DNA Technologies (IDT) codon optimization tool (www.IDTDNA.com). Plasmid pPTS422 carries the left TALE targeting domain, the 106 amino acids of *Burkholderia cenocepacia* DddA N-terminus and uracil-DNA glycosylase inhibitor (UGI). Transgene expression was driven by the maize ubiquitin 1 (Ubi1) promoter and nopaline synthase (nos) 3’-termination sequence (Tnos). Plasmid pPTS423 carries the right TALE targeting domain, the 30 amino acids DddA C-terminus and UGI. The transgene is driven by the rice polyubiquitin gene RUBQ2 promoter. Both TALE-DddA-UGI coding regions are fused in frame at their N-terminus to the 49 amino acids chloroplast targeting peptide (CTP) from maize Rubisco small subunit 2 (RbcS2). pPTS423 also carries a bar selectable marker gene cassette expressed from an enhanced Cauliflower Mosaic Virus (2xCaMV) 35S promoter and the CaMV polyA signal. The left TALE-DddA-UGI transgene cassette was cloned as a HindIII DNA fragment downstream and in the same orientation of the right TALE-DddA-UGI cassette to create the final pPTS424 vector.

For nuclear retransformation of the parent mutant lines, the chloroplast *rbcL* and *atpB* genes including the RbcS2 CTP were codon optimized for *Zea mays* nuclear expression. The transgenic *rbcL* gene carried a HiBit epitope tag and the *atpB* gene carried a 3X HA epitope tag at the C-terminus to distinguish the recombinant proteins. The transgenes were expressed from a 3X viral enhanced^61^ (FMV/PCSV/MMV) maize Ubi1 promotor and the Tnos terminator. The transgene cassettes were then cloned into a T-DNA vector carrying a hygromycin resistance selectable marker gene driven by 2XCaMV and CaMV polyA signal, to create plasmids pPTS451 carrying the nuclear *atpB* transgene and pPTS454 carrying the nuclear *rbcL* transgene.

### Generation of nonphotosynthetic chloroplast mutant lines

Plasmid DNAs were introduced into *Agrobacterium* strains LBA4404 Thy-^62^ or AGL1 (Intact Genomics, St. Louis MO) via electroporation using a Biorad MicroPulser. Selection for bacterial transformants was via resistance to 100 mg/L kanamycin, 50 mg/ L gentamycin, 50 mg/L Thymidine for LBA4404 Thy- or 50 mg/L Kanamycin and 15 mg/L rifampicin for AGL1. Integrity of intact plasmids was confirmed by colony PCR sequencing.

A wild-type maize hybrid genotype (Pa91 x H99) x H99 or a homozygous nuclear transgenic line overexpressing BBM and WUS2 morphogenic genes in an H99 background (Venkata RamanaRao Mangu and Murug Mookkan, manuscript in preparation) were used as recipients for *Agrobacterium* transformation using the AGL1 or LBA4404 Thy-strains, respectively. 10-12 days after pollination (DAP) immature embryos were transformed with Agrobacterium strains as described^47^ with minor modifications. Immature embryos were cocultivated in liquid MS media^63^ for 15 minutes, and then placed scutellum side up on solid ½ MS media for 2 days at 21°C in the dark. Embryos were then transferred onto a modified somatic embryo induction medium^44^ (SEIM) containing 0.7 g/L L-proline, no casein hydrolysate, and 250 mg/L carbenicillin (for LBA4404-Thy-) or 250 mg/L cefotaxime (AGL1). After 3-5 days resting at 28°C in the dark, the immature embryos were moved to SEIM for ∼8 weeks including 5 mg/L bialaphos (Gold Biotechnology, Inc. USA) for selection. Bialaphos resistant calli were moved onto maturation medium^44^ (MM) containing 10 mg/L phosphinothricin and incubated under dark conditions for 2 weeks or until signs of shoot regeneration. Phosphinothricin selection was maintained during the shoot regeneration phase to minimize escapes. Once shoots elongated to 3-4 cm length, the plantlets were moved onto MGM rooting medium^44^ under cool white fluorescent lights (40-60 umol) at 28°C with a 16/8 hr light/dark cycle. Candidate chloroplast mutant lines were identified as pale green or yellow shoots after 2-3 weeks in the light. Nuclear transgenesis was confirmed by PCR and PCR-sequencing detection of transgenic gene sequences. Chloroplast mutant lines were characterized as described in the text. All the media components were purchased from PhytoTech Labs, Inc. USA except where noted.

### Nuclear retransformation of chloroplast mutant lines

Embryogenic callus was established from leaf base cuttings dissected from regenerated homoplasmic chloroplast mutant lines and maintained on SEIM media in the dark. Freshly subcultured callus was precultured for 2-4 days on SEIM media, then transferred into the center of solid SEIM osmotic medium plates (SEIM modified to include 20 gm/L sucrose, 36.44 gm/L mannitol and 36.44 gm/L sorbitol) and cultured for an additional 2-4 hours in preparation for particle bombardment. The plasmid DNAs for pPTS454 and pPTS455 were coprecipitated onto gold particles in 1:1 w/w ratio. Particle bombardment was carried out as described.^64^ After an overnight resting period in the dark at 28°C, somatic embryos were placed on SEIM with 150 mg/L hygromycin for selection in the dark for 6-8 weeks. Hygromycin resistance callus was transferred onto MM with 50 mg/L hygromycin for an additional 2-3 weeks to regenerate plants. Green regenerated plantlets were moved onto MGM rooting medium as above. The presence of plasmid transgenes was confirmed by PCR-sequencing as described in the text.

### PCR-based Genotyping

Genomic DNA was extracted using the DNeasy Plant Mini Kit (Qiagen, CA, USA) and quantified using Qubit fluorometer (Thermo Fisher Scientific, MA, USA) according to the manufacturer’s instructions. Specific primers were designed to amplify regions flanking the gene-editing target sites in the chloroplast genome (Primer3, Snapgene Software, from Dotmatics; available at snapgene.com). Primer sequences for *Zea mays* genes tested are listed in Supplementary Table 4. PCR amplification of the targeted genomic regions was carried out using a thermal cycler (C1000, Bio-Rad Laboratories Inc., CA, USA). The reaction was performed in 15 μL volume containing 10 μL 2x Platinum SuperFi II PCR Master Mixes (Thermo Fisher Scientific, MA, USA), 500 nM of each primer and 10 ng of DNA. The PCR products were purified using a gel extraction kit (Zymoclean Gel DNA Recovery Kit, Zymo Research, CA, USA) and subsequently sequenced using Oxford Nanopore technology (Plasmidsaurus, KY, USA) to confirm the presence of the intended genetic modifications. The sequencing data were analyzed using Snapgene software to compare the amplified sequences against the reference genome.

### Chloroplast genome sequencing and analysis

All plant DNA extraction from frozen tissues, library preparation, and genome sequencing was performed by Novogene using an Illumina sequencing platform. Briefly, 0.2 μg DNA per sample was used as input material for the DNA library preparations. Sequencing library was generated using NEBNext® UltraTM DNA Library Prep Kit for Illumina (NEB, USA, Catalog #: E7370L) following manufacturer’s recommendations and index codes were added to each sample. Genomic DNA sample was fragmented by sonication to a size of 350 bp. Then DNA fragments were endpolished, A-tailed, and ligated with the full-length adapter for Illumina sequencing, followed by further PCR amplification. After PCR, products were purified by AMPure XP system (Beverly, USA). Subsequently, library quality was assessed on the Agilent 5400 system (Agilent, USA) and quantified by QPCR (1.5 nM). The qualified libraries were pooled and sequenced on Illumina platforms with PE150 strategy, according to effective library concentration and data amount required.

The original fluorescence image files obtained from DNBSEQ-T7 platform are transformed to short reads (Raw data) by base calling and these short reads are recorded in FASTQ^65^ format, which contains sequence information and corresponding sequencing quality information. After data processing, sequencing data was mapped to the reference genome using the Burrows-Wheeler Aligner^66^ to get the original mapping results stored in BAM format. Then, the SNP/InDel sets were called by SAMtools, using the following criteria: depth of the variate position >4, mapping quality >20. In some cases, sequence files were analyzed using Geneious software platform, that utilizes the same algorithms in a graphical user interface. The chloroplast reference genome used by Novogene for sequence mapping was *Zea mays* isolate Zhengdan 958 (Genbank MK348606.1) determined to be identical to our H99 chloroplast genome sequence based on our in-house de novo chloroplast genome assembly via the Geneious platform. SNPs were visualized on the Geneious software platform or the publicly available IGV viewer^67^ (www.igv.org).

### Protein extraction and gel blotting

Tissue was stored at −80°C. Tissue was ground to a powder in liquid nitrogen and protein extracted via the Sigma Plant Total Protein Extraction Kit, Mini (Sigma-Aldrich, MO, USA) following the manufacturer’s protocol with two changes; additional methanol incubations were performed until tissue was no longer green, and the protease inhibitor used was Plant ProteaseArrest (G-Biosciences). Proteins were quantified using the Qubit Broad Range Protein Assay (ThermoFisher Scientific, MA, USA).

Total protein was mixed with Novex Tris-Glycine SDS Sample Buffer (ThermoFisher Scientific) with 100 mM dithiothreitol, and then heated at 85°C for two minutes. Plant proteins and PageRuler Plus Prestained Protein Ladder (ThermoFisher Scientific) were loaded onto 4-20% Novex WedgeWell Tris-Glycine SDS PAGE gels (ThermoFisher Scientific). Running buffer was 1X Tris-Glycine SDS Buffer (VWR International, PA, USA). Gels were run at 225V, constant V, for 35 to 40 minutes. Protein was transferred from the gels to low fluorescence PVDF membranes (Azure Biosystems, CA, USA) in 1X Azure Transfer Buffer (Azure Biosystems). The transfer was run at 300mA, constant A, for 60 minutes. Membranes were stained with Ponceau S (Sigma-Aldrich) to observe relative protein loading.

For protein gel blots we used the AzureSpectra Fluorescent Western Blotting Kit with Fluorescent Block (Azure Biosystems) following the manufacturer’s protocol. For HiBit detection, we first performed the fluorescent antibody western and then followed with the Nano-Glo HiBit Blotting System (Promega, WI, USA) following the manufacturer’s protocol for PVDF membranes. The antibodies were: rabbit anti-atpB 1:1000 (PhytoAB, PHY0312;); rabbit anti-rbcL 1:1000 (PhytoAB, PHY0396); mouse anti-HA 1:1000 (Genscript, A01296); goat anti-rabbit-650 1:2500 (Azure Biosystems); and goat anti-mouse-550 1:2500 (Azure Biosystems).

An Azure 400 Imager (Azure Biosystems) was used to image the blots. For fluorescent blots the default settings for Cy3 and Cy5 dyes were used. For HiBit detection, the chemiluminescence channel was used in cumulative mode for 30 minutes (10 × 3 minutes). ImageJ software was used to separate channels, convert to grayscale, and invert Images. Original protein gel image files are available in the Supplemental Materials.

### Measurements of photosynthetic parameters

High-resolution fluorescence images were taken using CropReporter (PhenoVation, Wageningen, The Netherlands). False-color images were generated by the PhenoVation Data Analysis software. *F_v_/F_m_* measurements were made using an OJIP protocol.^68^ A modification to the Pulse Amplitude Modulation assay was made to measure NPQ following a light induction curve.^69^ Briefly, cultures were grown under identical light and temperature conditions (50-60 μmol photons m^-2^ · s^-1^; 28 °C) until at plantlets with at least two true leaves had emerged. Immediately before the assay, plantlets were dark adapted for 20 minutes, and then subjected to saturating pulses of actinic light (185 μmol photons m^-2^ · s^-1^).

## Acknowledgements

The authors would like to thank members of the Plastomics team for ongoing support during all aspects of this work and for a critical reading of the manuscript by Charles Armstrong. We especially want to thank Murug Mookkan for early work in helping to create the BBM/WUS line used here, and Bill Gordon-Kamm for providing materials. We also thank the Danforth Plant Science Center and Sally Fabbri, Dan Long and Katrina Wilsey for greenhouse support, the Danforth Phenotyping Core Facility, and Noah Fahlgren help with the Crop Reporter data. Some aspects of this work were funded in part by an NSF SBIR grant (2015099) to JMS. JMS thanks Riqing Li, Si Nian Char and Bing Yang for cloning of a precursor plasmid.

---

**Supplemental Figure 1.**
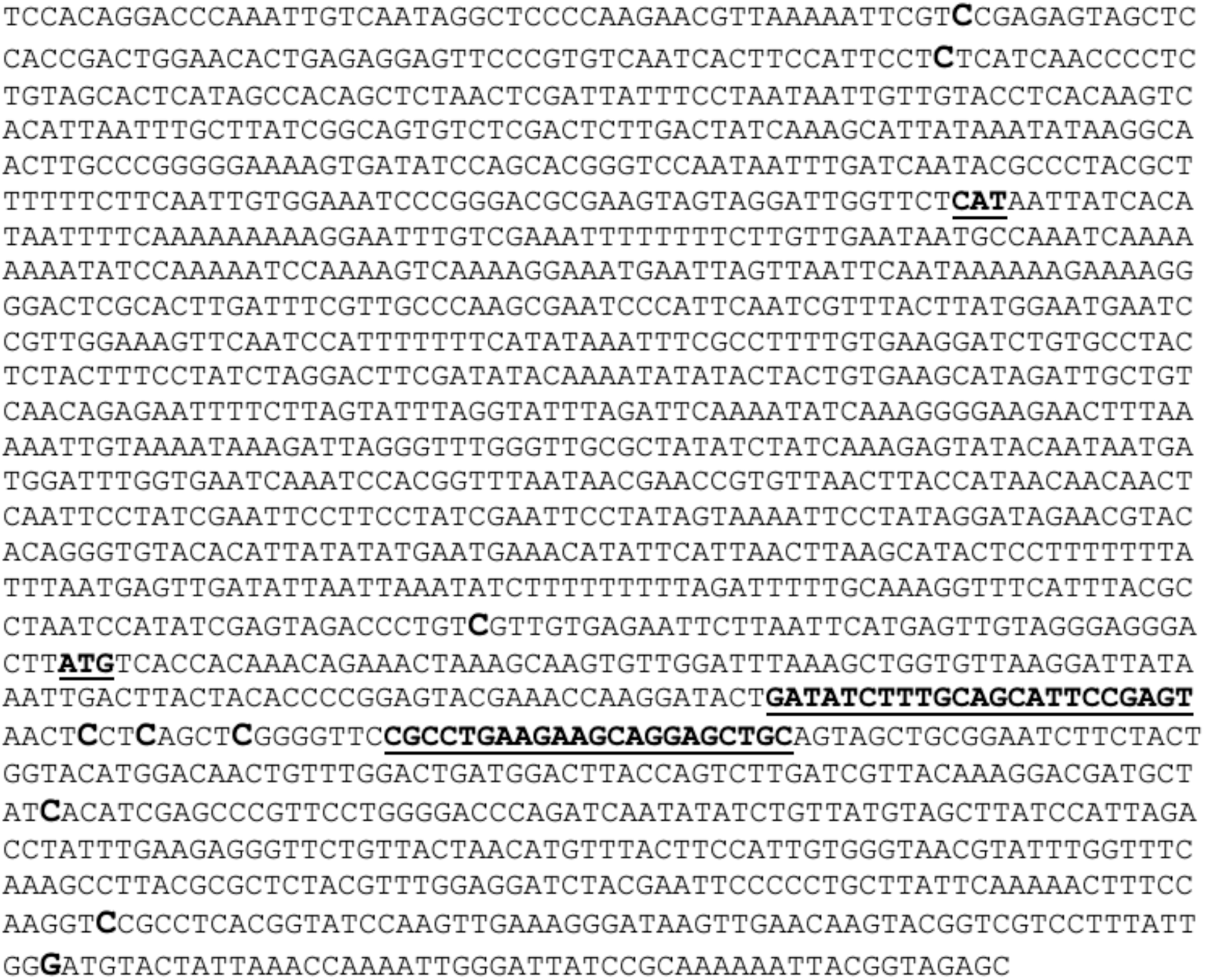
Sequence of the *rbcL – atpB* edited chloroplast gene regions. The start of the *rbcL* coding region is denoted by the ATG start codon (bold underlined) and the start of the divergent *atpB* coding region is denoted by its CAT start codon (bold underlined, opposite DNA strand). The two coding regions are separated by 781 bp. The TALE left and TALE right binding sites (bold, underlined) in the *rbcL* coding region are shown along with the three cytosine residues (bold, larger font) that are potential targets for editing. Observed off-target cytosine or guanosine on the opposite DNA strand are also shown in bold.

**Supplementary Figure 2.**
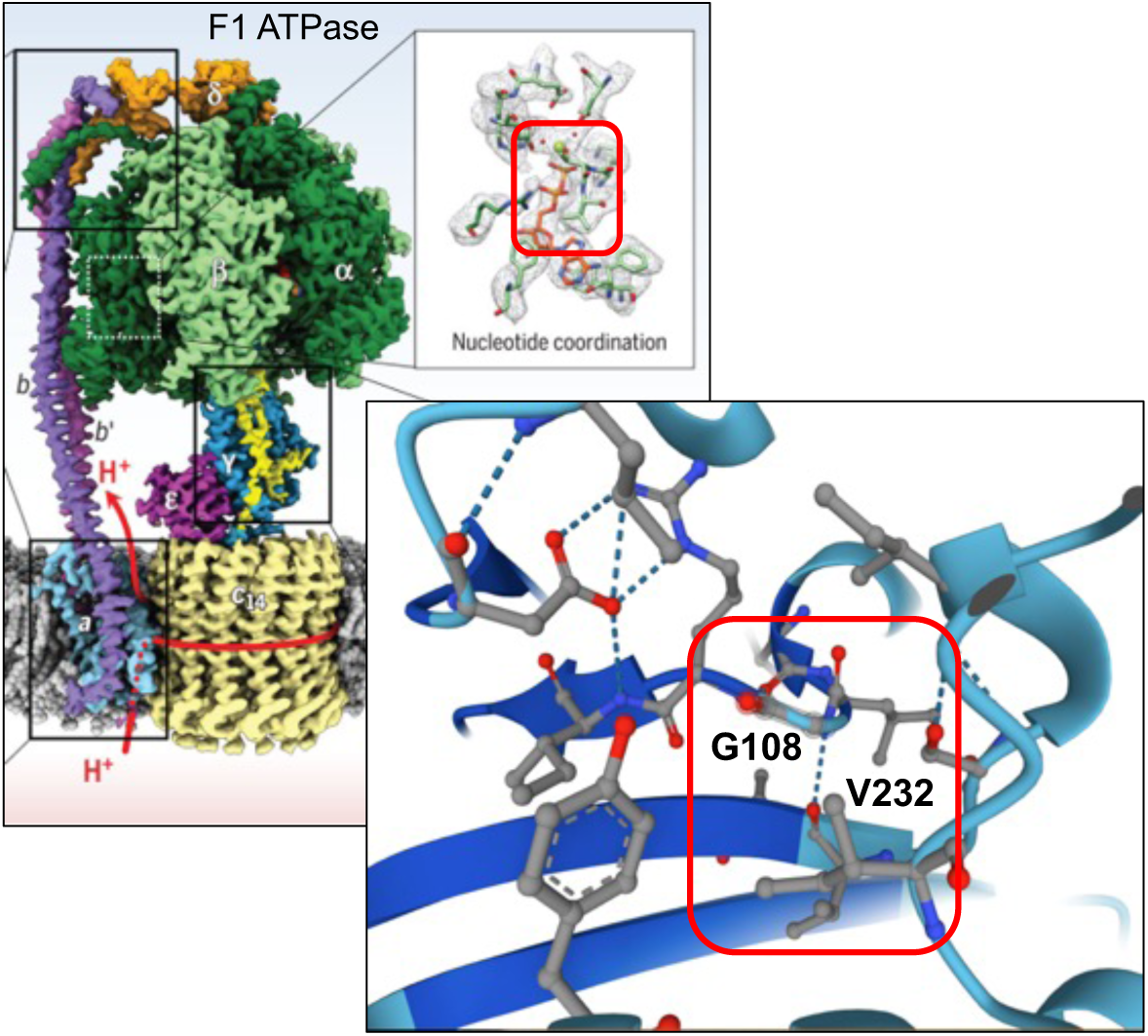
Structure of the N-terminal nucleotide binding region of the AtpB protein within the F1 ATPase complex. Alphafold software (www.alphafold.com) was used to model the N-terminal region of the AtpB protein coding region. Amino acid glycine 108 (G108) is coordinated (shown by red box) with valine 232 (V232), suggesting that the G108E mutation must disrupt the three-dimensional structure and binding of ADP+ and *Pi* to the F1 ATPase complex. ATP Synthase structure model modified from Hahn et al.^55^

**Supplementary Figure 3.**
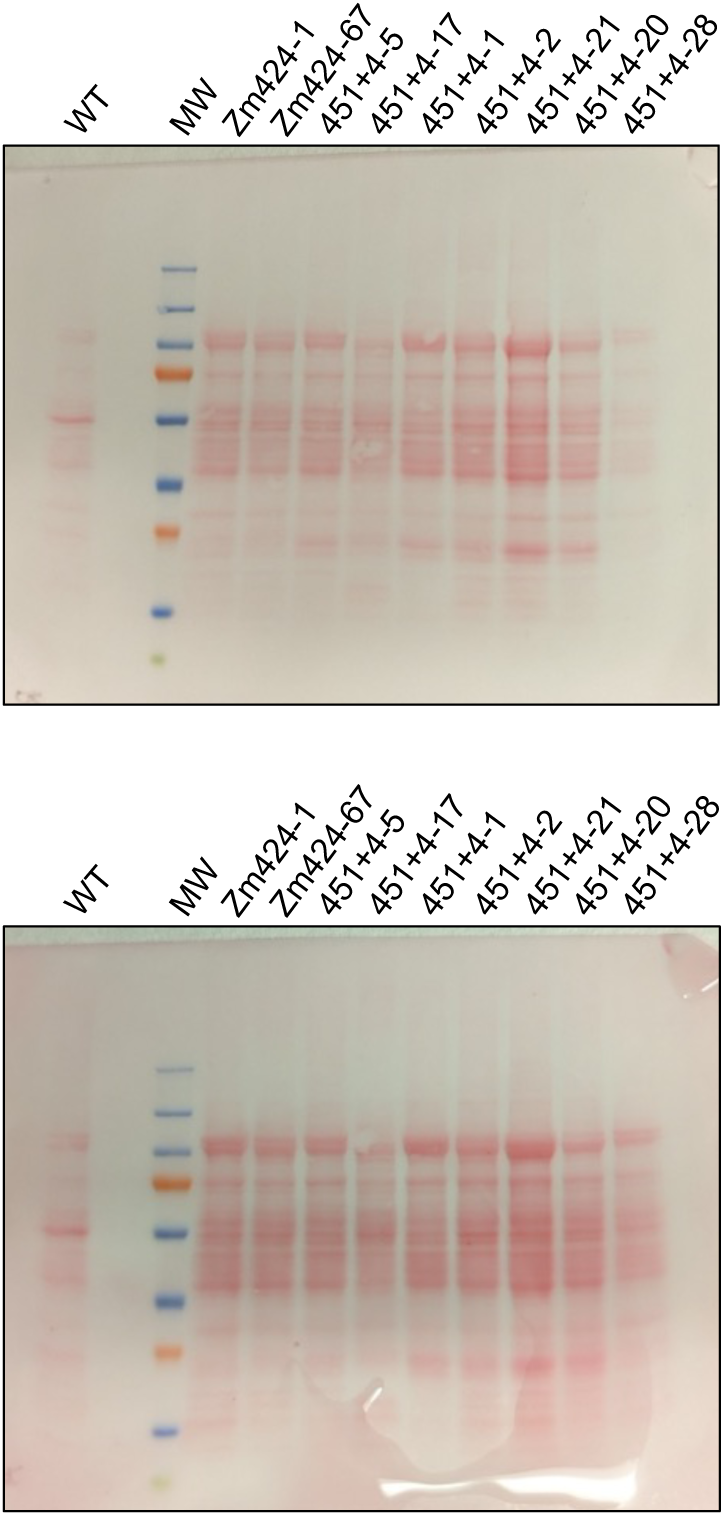
Ponceau staining of protein gel blots shown in Figure 3. (A) Protein gel blot used to hybridize to AtpB protein and HA epitope tag antibodies. (B) Protein gel blot used to hybridize with RbcL protein antibody and to detect luminescence via the HiBit epitope tag.

**Supplementary Figure 4.**
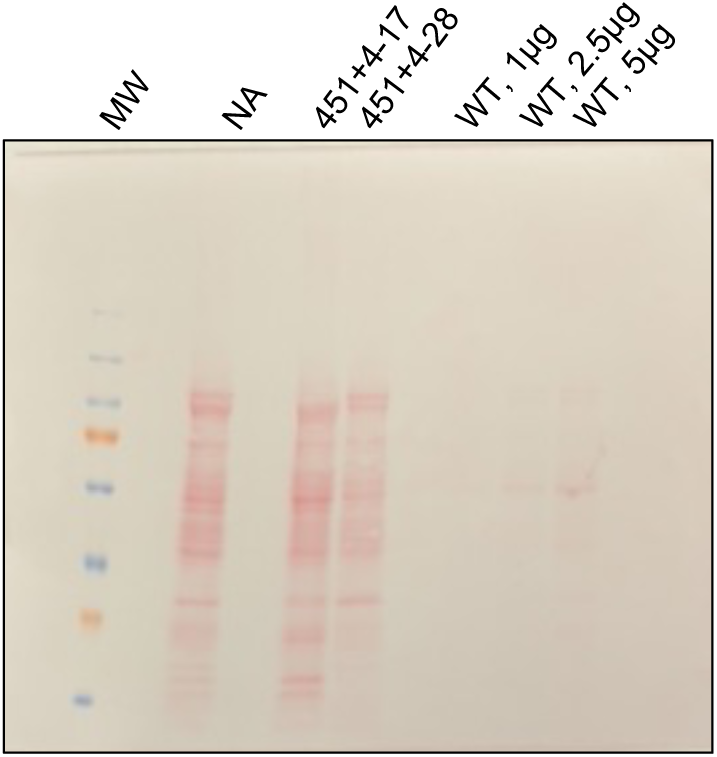
Ponceau staining of protein gel blots shown in Figure 4. (A) Protein gel blot used to hybridize to AtpB protein and HA epitope tag antibodies, used to estimate recombinant AtpB protein levels in nuclear transgenic lines compared to wild-type plants.

**Supplementary Table I.**
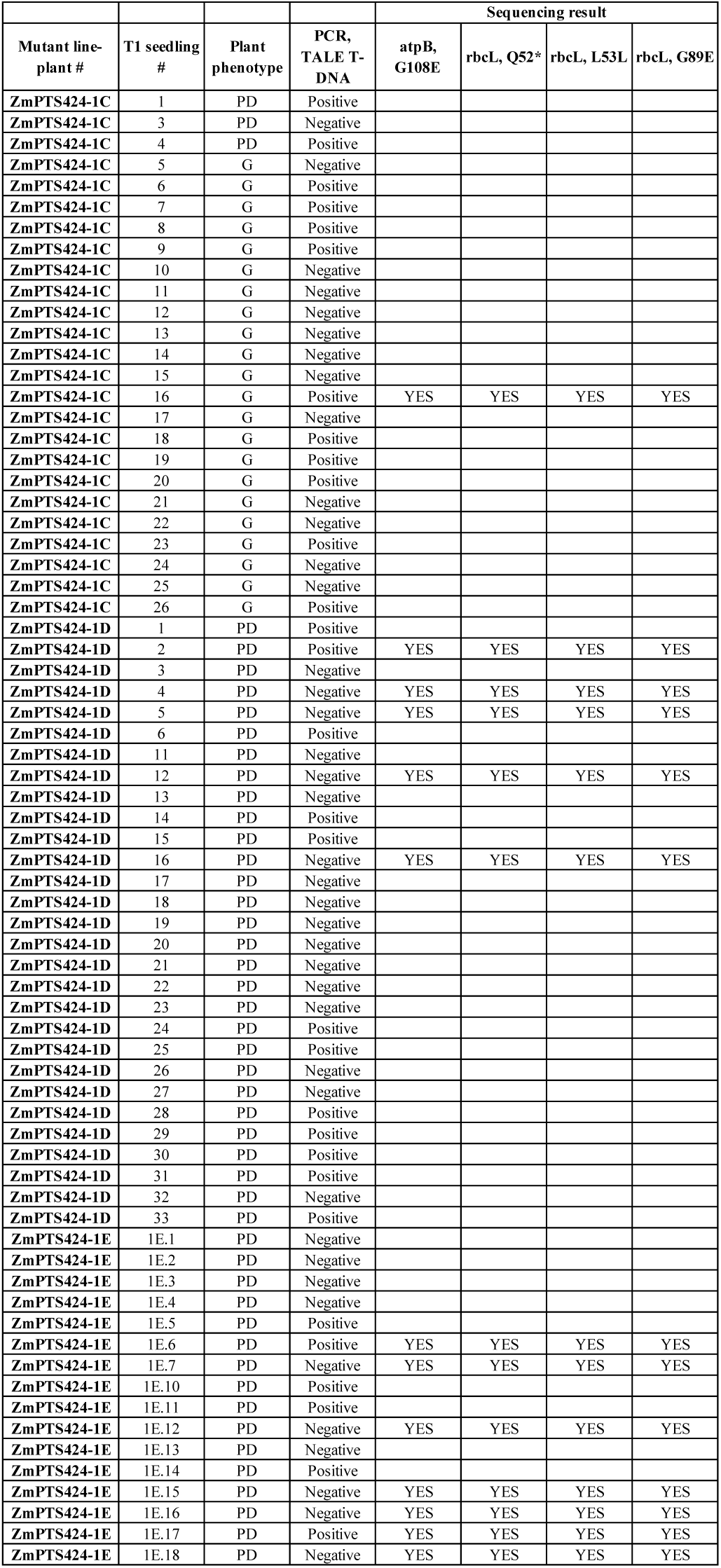
Segregation of the nuclear TALE-DddA-UGI locus in T1 seedlings derived from chimeric mutant ZmPTS424-1 plants grown in the greenhouse and crossed as females with wild-type pollen. ZmPTS424-1C, 1D and 1E refer to independently regenerated chimeric mutant plants that were grown to maturity in the greenhouse. Numerous seedlings were sown in soil and their phenotypes recorded as pigment deficient (PD) or green (G). PCR analysis was used to determine if the TALE-DddA-UGI locus is present in the T1 seedlings and PCR-sequencing was used to confirm the presence of the original chloroplast mutations in a subset of seedlings.

**Supplementary Table 2.** Phenotype and genotypes of nuclear re-transformed lines after co-transformation of the parental mutants with vectors pPTS451 and pPTS454. Independently re-transformed lines (451+4 event ID represented by a number) were regenerated in some cases into multiple plants (represented by lower case letters). The phenotype of each regenerated plant was recorded as green (G) or pigment deficient (PD). PCR analysis was used to confirm the presence of the nuclear transgenic *CTP-atpB*, *CTP-rbcL* or hygromycin resistance (*Hyg*) genes. NT, not tested. Note that the presence of the *CTP-rbcL* gene is independent of whether the lines are green or pigment-deficient, as there are no cases of a green phenotype exclusively testing PCR-positive for the *CTP-rbcL* transgene. This indicates that the greening phenotype is associated with the *CTP-atpB* gene. In some instances, however, green lines tested PCR-negative for the *CTP-atpB* gene, which could be attributed to false-negative results or structural alterations in the transgenic sequence.

| Parent mutant line | 451+4 event ID | Phenotype | PCR, *CTP-aptB* | PCR, *CTP-rbcL* | PCR, *Hyg* |
| --- | --- | --- | --- | --- | --- |
| ZmPTS424-67 | 25a.1 | G | Positive | Positive | Positive |
| ZmPTS424-67 | 25a.2 | G | Positive | Positive | Positive |
| ZmPTS424-67 | 25a.3 | G | Positive | Positive | Positive |
| ZmPTS424-67 | 25a.4 | G | Positive | Negative | Positive |
| ZmPTS424-67 | 17a.5 | G | Negative | Negative | Positive |
| ZmPTS424-67 | 17a.6 | G | Positive | Positive | Positive |
| ZmPTS424-67 | 17a.7 | G | Positive | Negative | Positive |
| ZmPTS424-67 | 17a.8 | G | Negative | Negative | Positive |
| ZmPTS424-67 | 17a.9 | G | Negative | Negative | NT |
| ZmPTS424-67 | 17a.10 | G | Positive | Positive | NT |
| ZmPTS424-67 | 21b.11 | G | Positive | Positive | NT |
| ZmPTS424-67 | 21b.13 | G | Positive | Positive | NT |
| ZmPTS424-67 | 21b.14 | G | Positive | Positive | NT |
| ZmPTS424-67 | 21b.15 | G | Negative | Negative | NT |
| ZmPTS424-67 | 21b.16 | G | Positive | Positive | NT |
| ZmPTS424-67 | 17a.17 | PD | Negative | Positive | NT |
| ZmPTS424-67 | 17a.18 | PD | Negative | Positive | NT |
| ZmPTS424-67 | 17a.19 | PD | Negative | Positive | NT |
| ZmPTS424-67 | 17a.20 | PD | Negative | Positive | NT |
| ZmPTS424-67 | 17a.21 | PD | Negative | Positive | NT |
| ZmPTS424-67 | 17a.22 | PD | Negative | Positive | NT |
| ZmPTS424-67 | 17b | G | Positive | Positive | NT |
| ZmPTS424-67 | 20e | G | Positive | Positive | NT |
| ZmPTS424-67 | 20f | G | Positive | Positive | NT |
| ZmPTS424-67 | 20g | G | Positive | Positive | NT |
| ZmPTS424-67 | 20h | G | Positive | Positive | NT |
| ZmPTS424-67 | 20i | G | Positive | Positive | NT |
| ZmPTS424-67 | 20j | G | Positive | Positive | NT |
| ZmPTS424-67 | 20k | G | Positive | Positive | NT |
| ZmPTS424-67 | 20n | G | Positive | Positive | NT |
| ZmPTS424-67 | 20p | G | Positive | Negative | NT |
| ZmPTS424-67 | 20q | G | Positive | Positive | NT |
| ZmPTS424-67 | 21e | G | Positive | Positive | NT |
| ZmPTS424-67 | 22e | G | Positive | Positive | NT |
| ZmPTS424-67 | 22f | G | Positive | Positive | NT |
| ZmPTS424-67 | 22g | G | Positive | Positive | NT |
| ZmPTS424-67 | 22h | G | Positive | Positive | NT |
| ZmPTS424-67 | 22i | G | Positive | Positive | NT |
| ZmPTS424-67 | 22j | G | Positive | Positive | NT |
| ZmPTS424-67 | 22k | G | Positive | Positive | NT |
| ZmPTS424-67 | 22l | G | Positive | Positive | NT |
| ZmPTS424-67 | 22m | G | Positive | Positive | NT |
| ZmPTS424-67 | 22n | G | Positive | Positive | NT |
| ZmPTS424-67 | 22p | G | Positive | Negative | NT |
| ZmPTS424-67 | 22q | G | Positive | Negative | NT |
| ZmPTS424-67 | 22r | G | Positive | Negative | NT |
| ZmPTS424-67 | 23c | G | Positive | Negative | NT |
| ZmPTS424-1 | 1A | G | Positive | Positive | NT |
| ZmPTS424-1 | 1b | G | Positive | Negative | NT |
| ZmPTS424-1 | 1c | G | Positive | Negative | NT |
| ZmPTS424-1 | 1d | G | Positive | Negative | NT |
| ZmPTS424-1 | 4 | G | Positive | Positive | NT |
| ZmPTS424-1 | 5a | G | Positive | Positive | NT |
| ZmPTS424-1 | 5b | G | Positive | Positive | NT |
| ZmPTS424-1 | 9a | PD | Positive | Positive | NT |
| ZmPTS424-1 | 9b | PD | Positive | Positive | NT |
| ZmPTS424-1 | 9c | PD | Positive | Negative | NT |
| ZmPTS424-1 | 9d | PD | Negative | Positive | NT |
| ZmPTS424-1 | 10a | PD | Negative | Negative | Positive |
| ZmPTS424-1 | 10b | PD | Negative | Negative | Positive |
| ZmPTS424-1 | 10c | PD | Negative | Negative | Positive |
| ZmPTS424-1 | 10d | PD | Negative | Negative | Positive |
| ZmPTS424-1 | 13 | PD | Negative | Positive | NT |
| ZmPTS424-1 | 14a | PD | Negative | Positive | NT |
| ZmPTS424-1 | 14b | PD | Negative | Negative | Positive |
| ZmPTS424-1 | 15 | PD | Negative | Positive | NT |
| ZmPTS424-1 | 1a | G | Positive | Negative | NT |
| ZmPTS424-1 | 1c | G | Positive | Negative | NT |
| ZmPTS424-1 | 1d | G | Negative | Negative | Positive |
| ZmPTS424-1 | 9a | PD | Negative | Negative | Positive |
| ZmPTS424-1 | 9b | PD | Negative | Negative | Positive |
| ZmPTS424-1 | 9c | PD | Negative | Negative | Positive |
| ZmPTS424-1 | 9d | PD | Negative | Positive | NT |
| ZmPTS424-1 | 14a | PD | Negative | Negative | Positive |
| ZmPTS424-1 | 14b | PD | Negative | Negative | Positive |

**Supplementary Table 3.** Summary of *Agrobacterium-*mediated transformation of vector pPTS424 into immature embryos from the BBM/WUS2 line (in a H99 genetic background) and wild-type genotype (Pa91xH99) x H99. The number of T0 bialaphos resistant events is shown along with the number that had a pigment deficient phenotype. Note that one pigment deficient line in the H99 (BBM/WUS) genetic background carried multiple chloroplast genome mutations outside of the *atpB* and *rbcL* genes, as was not characterized further.

| Genotype | Agrobacterium strain used for transformation | Number of immature embryos transfected | Number of T0 events selected on bialaphos | Number of independent pigment deficient transgenic events |
| --- | --- | --- | --- | --- |
| H99 (BBM/WUS2) | LBA4404 Thy- | 125 | 36 | 3 |
| (Pa91x H99) xH99 | AGL1 | 100 | 9 | 1 |

**Supplementary Table 4.**
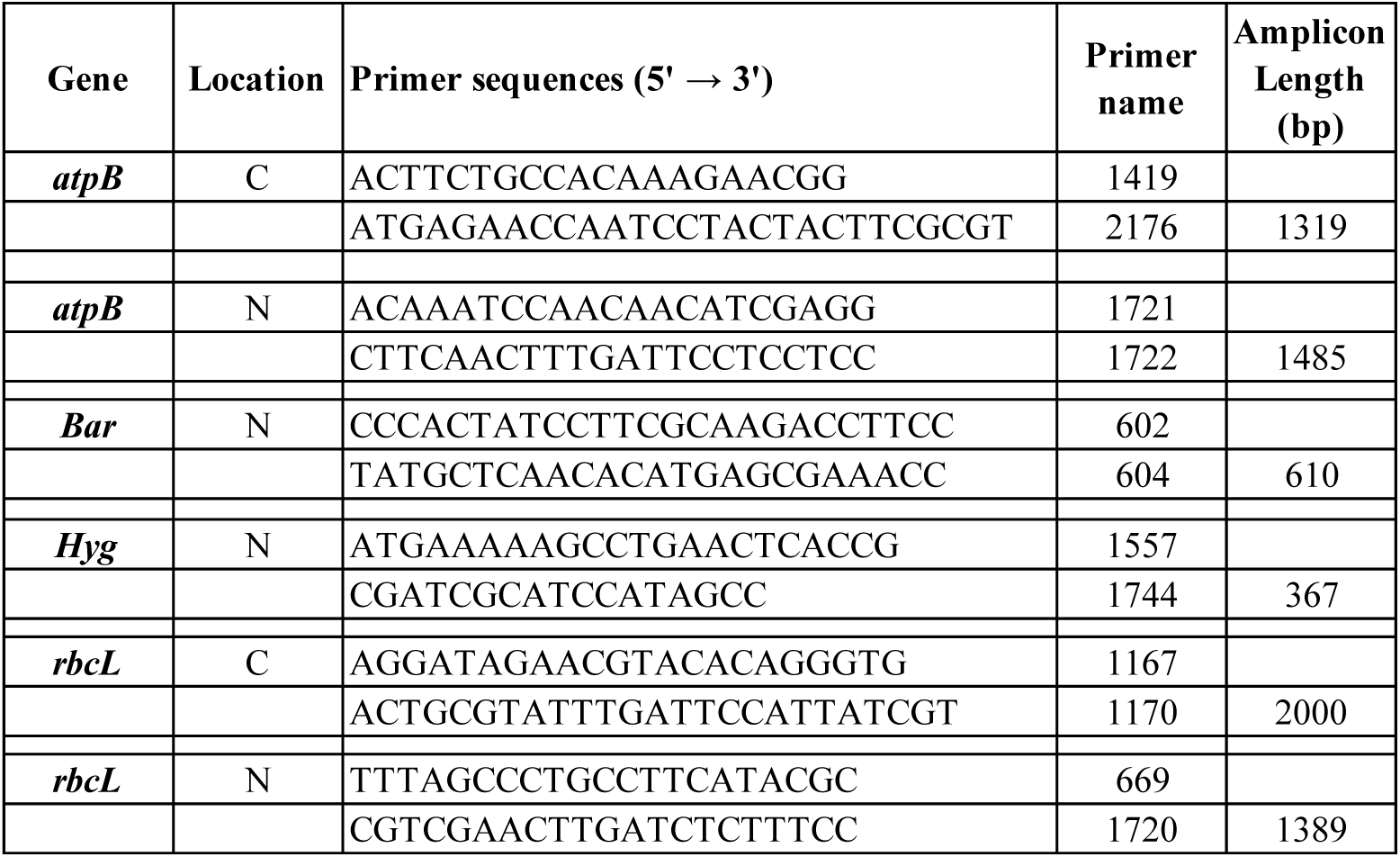
Oligonucleotide primers used in this report to amplify nuclear (N) transgenes and endogenous chloroplast (C) sequences.

